# APPLE-MS: A affinity purification-mass spectrometry method assisted by PafA-mediated proximity labeling

**DOI:** 10.1101/2025.05.06.652320

**Authors:** Shihan Luo, Lijuan Xie, Lin Yang, Zheyao Hu, Lei Wang, Yueqin Wang, Qingqing Li, Shujuan Guo, Shengce Tao, Hewei Jiang

## Abstract

While affinity purification-mass spectrometry (AP-MS) has significantly advanced protein-protein interaction (PPI) studies, its limitations in detecting weak, transient, and membrane-associated interactions remain. To address these challenges, we introduced an innovative proteomic method termed <u>A</u>ffinity <u>P</u>urification coupled <u>P</u>roximity <u>L</u>ab<u>E</u>ling-Mass Spectrometry (APPLE-MS), which combines the high specificity of Twin-Strep-tag enrichment with PafA-mediated proximity labeling. This method achieves unprecedented sensitivity while maintaining high specificity (4.07-fold over AP-MS). APPLE-MS also revealed the dynamic mitochondrial interactome of SARS-CoV-2 ORF9B during antiviral responses, while endogenous PIN1 profiling uncovered novel roles in DNA replication. Notably, APPLE-MS enabled *in situ* mapping of GLP-1 receptor complexes, demonstrating its unique capabilities for membrane PPI studies. This versatile method advances interactome research by providing comprehensive, physiologically relevant PPI networks, opening new opportunities for mechanistic discovery and therapeutic targeting.

**GRAPHICAL ABSTRACT:** 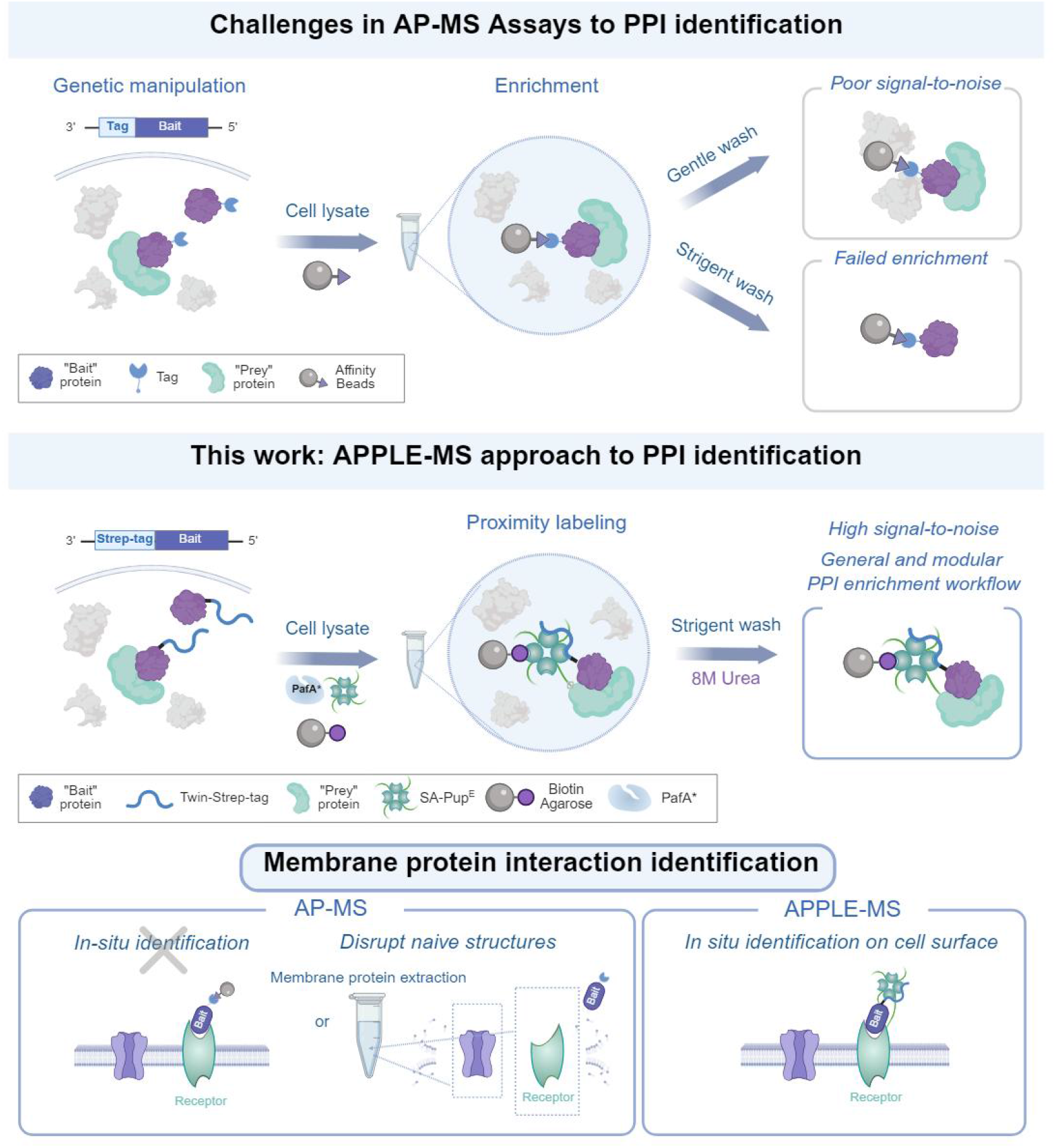

**MOTIVATION – *Cell Reports Methods* only:** Affinity purification-mass spectrometry (AP-MS) has become a widely used method for capturing high-affinity protein-protein interactions (PPIs). However, it still faces several limitations, including high levels of non-specific binding, challenges in detecting weak interactions, and limited ability to identify PPIs at the cell surface in situ. To overcome these challenges, we introduce a modular AP-MS method enhanced by PafA-mediated proximity labeling, which improves both the specificity and sensitivity of PPI detection in a single, streamlined workflow.

**SIGNIFICANCE – *Cell Chemical Biology* only:** Proximity labeling is a powerful tool in chemical biology, providing critical support for studying protein function, cell signaling, and disease mechanisms by chemically labeling and capturing biomolecular interactions. In this study, we introduce a modular affinity purification-mass spectrometry (AP-MS) method that integrates proximity labeling. This method leverages the proximity-dependent enzymatic activity of PafA to covalently attach Pup^E^ to nearby proteins, in combination with the high-affinity interaction between Twin-Strep-tag and streptavidin to achieve efficient binding. This innovative approach enables precise identification of protein-protein interactions (PPIs), offering a powerful tool for capturing weak and transient interactions that are often missed by traditional AP-MS techniques. By enhancing both the specificity and efficiency of protein labeling, our work provides a robust chemical biology strategy to elucidate complex protein interactions, which are essential for understanding cellular processes and developing targeted therapeutic strategies, offering new insights into protein function and interaction dynamics in both health and disease.

## INTRODUCTION

Protein–protein interactions (PPIs) underpin almost every biological pathway, with more than 80% of proteins functioning within complexes to execute their roles in cells^1^. Dissecting PPIs is therefore essential not only for deciphering molecular mechanisms but also for identifying potential therapeutic targets. However, PPIs are inherently complex and dynamic, influenced by post-translational modifications (PTMs), cellular context, and transient binding events^2^. Over the years, numerous methods have been developed to tackle these challenges, ranging from yeast two-hybrid (Y2H) screens to advanced computational predictions^3,4,5^. Despite this diversity, each approach faces technical constraints such as high false-positive rates, suboptimal detection sensitivity, or limited utility in detecting weak or transient interactions.

Affinity purification–mass spectrometry (AP–MS) is one of the most widely adopted methods for defining PPIs in biology laboratories, owning to its established protocols and broad applicability, and its integration with quantitative proteomics, AP–MS has facilitated large-scale network studies^6^. However, a persistent challenge lies in the high sensitivity to operational details, small variations in execution often lead to inconsistent results among different operators. Additionally, the risk of non-specific binding remains a significant drawback. Tandem affinity purification (TAP) partially addresses this issue by introducing sequential purification steps; however, the stringent washes involved can inadvertently remove weak interactions, creating a trade-off between sensitivity and specificity^7^. AP–MS has also been used to analyze cell-surface PPIs by introducing affinity tags prior to cell lysis or by detergent-based membrane protein extraction. However, pre-lysis labeling methods frequently induce artificial receptor clustering through harsh washing conditions(e.g., low PH elution)^8^, while detergent-based extraction protocols disrupt critical membrane microdomains and lead to significant loss of peripheral interactors. For example, LPG14 — a commonly used anionic detergent — has been shown to dissociate up to 75% of interacting partners from NBD1, underscoring the inherent compromise between effective solubilization and maintaining native interactions^9^. These technical constraints lead to significant underrepresentation of biologically relevant but labile interactions, particularly transient complexes (t_1/2_ < 2 min) and low-affinity assemblies (K_d_ >100 μM) that dominate cell signaling processes. Proximity labeling–mass spectrometry (PL–MS) offers a complementary route to map protein networks^10^. By harnessing engineered enzymes or chemical probes that label proximal proteins, PL–MS excels at capturing weak or transient associations, facilitating the discovery of novel PPIs. In workflows such as BioID or APEX, affinity purification with streptavidin beads is integral for enriching the labeled proteins, enabling high-confidence identification via subsequent MS. Although sophisticated strategies combining BioID and AP–MS into a single workflow have been reported^11^, a major caveat is the need to fuse the labeling enzyme (typically 20–50 kDa) to the protein of interest. The introduction of additional mass may disrupt proper protein folding or molecular interactions, and the insertion of such an enzyme at the endogenous locus poses significant technical challenges compared to endogenous AP-MS, which requires only a small epitope tag.

Here, we introduce <u>A</u>ffinity <u>P</u>urification coupled <u>P</u>roximity <u>L</u>ab<u>E</u>ling–<u>M</u>ass <u>S</u>pectrometry (APPLE–MS), a proteomic interactome profiling method that combines the convenience of AP–MS with the covalent capture of transient interactions afforded by proximity labeling. By exploiting the proximity-dependent enzymatic activity of PafA to label neighboring proteins covalently with Pup^E^, our approach captures PPIs in their native cellular context. The strong binding affinity between the Twin-Strep-tag and streptavidin facilitates efficient enrichment and subsequent MS analysis. This design yields high sensitivity for detecting low-abundance or transient interactions and, crucially, permits *in situ* identification of cell-surface PPIs. We validated APPLE–MS by systematically interrogating intracellular and extracellular PPIs, highlighting its ease of operation, adaptability, and reliability in uncovering interaction networks across distinct cellular compartments.

## RESULTS

### APPLE enables the detection of PPIs

PafA, a central enzyme in the prokaryotic ubiquitin-like protein (Pup) tagging system, has been demonstrated in multiple studies to exhibit proximity labeling activity by catalyzing the ATP-dependent covalent attachment of the C-terminal glutamate residue of Pup^E^ to lysine side chains on target proteins^12^. Multiple studies have demonstrated that PafA exhibits robust proximity labeling activity, enabling the covalent ligation of Pup^E^ to lysine residues on proximal interactors. Leveraging this proven proximity labeling capability and the high affinity between Twin-Strep-tag and streptavidin^13^, we designed APPLE-MS to identify PPIs (Figure 1A). In this approach, a Twin-Strep-tag is fused to the bait protein, enabling its efficient capture by streptavidin. Upon addition of PafA and SA-Pup^E^, Pup^E^ is covalently attached to neighboring proteins, facilitating subsequent mass spectrometry-based identification of interactors. This method isolates PPIs under stringent washing conditions through the high-affinity biotin–streptavidin interaction, allowing for efficient purification of prey proteins and the capture of PPIs in a single integrated workflow.

**Figure 1.**
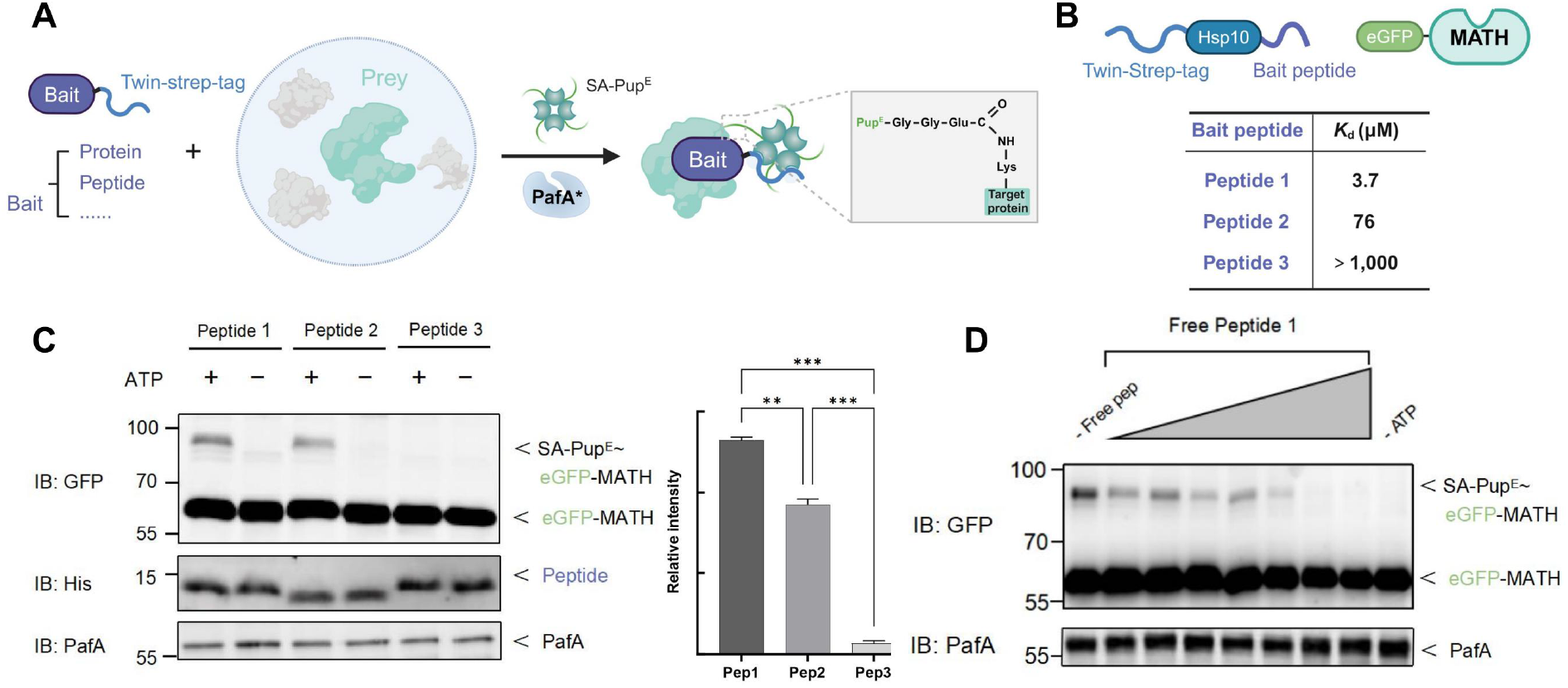
Design and validation of the APPLE proximity labeling system. (A)Schematic design of APPLE **proximity** labeling system. The bait protein (purple) is fused to a Twin-Strep-tag (blue) for streptavidin-Pup^E^ (SA-Pup^E^) recruitment. PafA mediates covalent pupylation of proximal lysine residue(s) on the prey protein(Green), converting the transient interaction between the bait and prey into stable SA-Pup^E^–prey conjugates. (B)Model system for evaluating the affinity detection limit in APPLE assay. Three Twin-Strep-tagged peptides (1-3) with MATH domain binding affinities ranging from 10^3^-10^6^ M^-1^ were designed. Binding affinity to the MATH domain decreases sequentially from peptide 1 to peptide 3. (C)**Quantitative validation of affinity detection limit**. APPLE assay with purified GFP-tagged MATH and affinity-varied peptides. Data represent mean±SEM from three independent experiments, ** P<0.01 and *** P<0.001 (two-tailed unpaired t-test) (D)**Competition binding assay**. Dose-dependent inhibition of Twin-Strep-tagged peptide 1 binding by untagged peptide 1 in the APPLE system.

To benchmark the APPLE-MS and empirically determine its detectable affinity range, we tested it against a well-characterized model system (Figure 1B). The dimeric MATH domain of SPOP recognizes small peptides containing distinct motifs via conserved hydrophobic and polar interactions, with sequence variations among substrates leading to differences in binding affinity^14^. Peptide 1 demonstrates the highest binding affinity for the MATH domain (3.7 μM), followed by peptide 2 (76 μM) and peptide 3 (>1000 μM), reflecting diminishing affinities due to alterations in their SBC motif sequences. We genetically fused the Twin-Strep-tag to each peptide and performed an APPLE-MS assay using the GFP-tagged MATH domain as the binding substrate. Our method successfully detected the SA-Pup^E^–MATH complex for peptide 1 and peptide 2 (Figure 1C), demonstrating the ability of APPLE-MS to capture weak interactions with affinities of at least 76 μM. Furthermore, increasing concentrations of free peptide 1 progressively inhibited MATH modification by outcompeting the Twin-Strep-tagged version in the APPLE-MS assay (Figure 1D). Altogether, these features enable APPLE-MS to robustly capture both stable and weak interactions.

### APPLE-MS to identify the ORF9b interactome

To further validate the applicability of APPLE-MS for systematic PPI screening, we employed this approach to comprehensively characterize the interactome of SARS-CoV-2 ORF9B. As a principal immune evasion factor, ORF9B suppresses host innate immunity by interacting with the mitochondrial translocase receptor TOM70, thereby inhibiting the recruitment of Hsp90 and associated signaling proteins, which impairs the interferon response^15,16^. However, its complete interactome remains poorly characterized, primarily due to limited sensitivity of existing methods (e.g., AP-MS) in detecting weak or transient interactions. These limitations prompted us to select ORF9B as an ideal model for mapping the APPLE-MS-based interactome, leveraging the method’s sensitivity to capture both stable immune regulators and weak viral-host interactions (Figure 2A).

**Figure 2.**
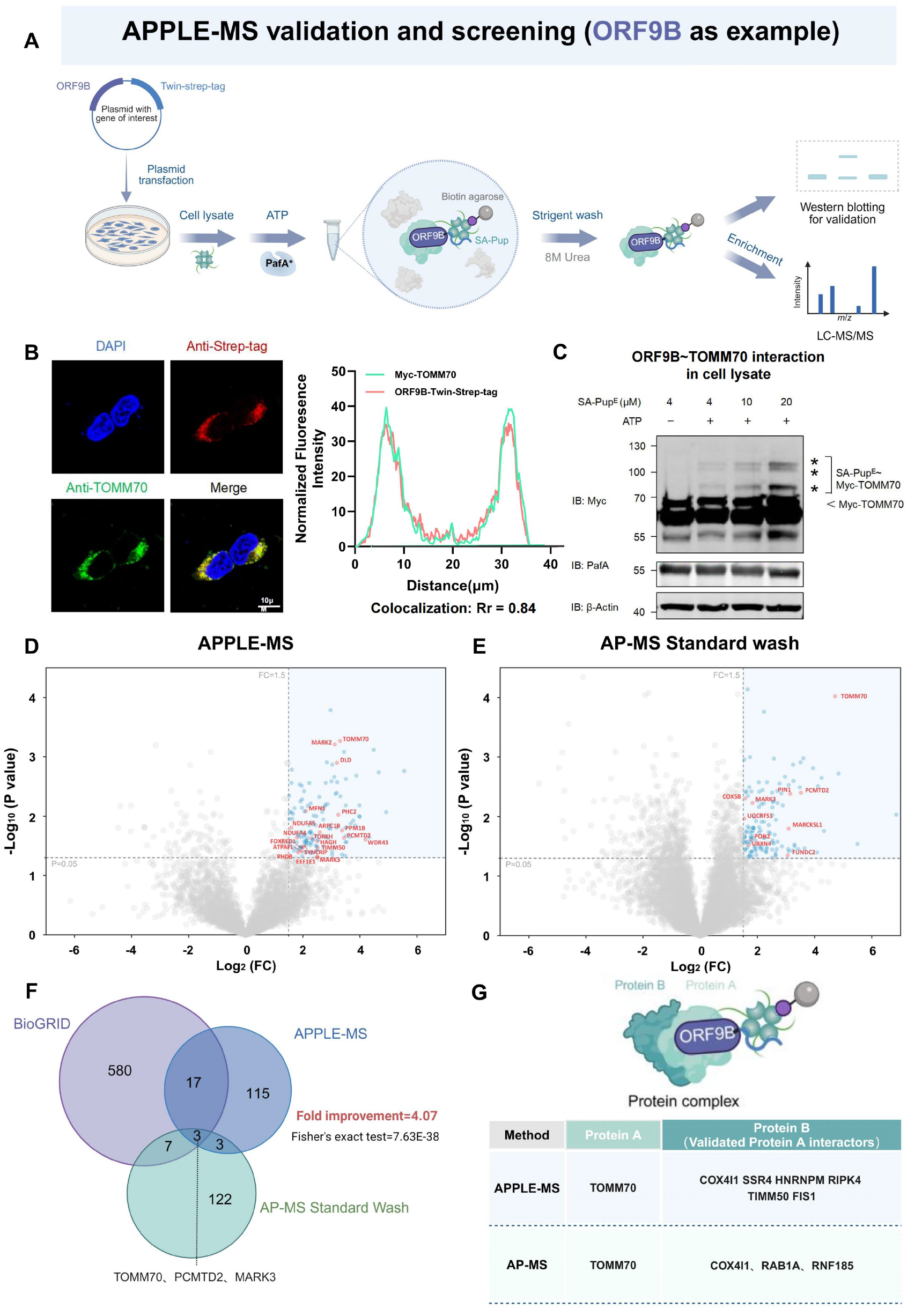
APPLE-MS to identify ORF9B interactome. (A)APPLE-MS **workflow**. Following pupylation of the prey protein, SA-Pup^E^-prey protein complexes are enriched using biotin agarose under stringent washing conditions, e.g., 8 M urea. Protein interactions are validated by SDS-PAGE or identified by label-free quantitative mass spectrometry.. (B)Subcellular localization of ORF9B. Confocal microscopy shows co-localization (yellow) of TOMM70 (green) and ORF9B-StrepTagII (red) in HEK293T cells. Nuclei were counterstained with DAPI (blue). Scale bars, 10 µm (representative of n=3). Normalized fluorescence intensity profiles along the spatial axis (distance) marked in the right panel. Quantification of Pearson’s correlation coefficient from three independent experiments. (C)Immunoblotting analysis of APPLE assay with HEK293T cell lysates co-expressing TOMM70 and ORF9B-Twin-Strep-tag. (D-E) ORF9B interactome profiling by APPLE-MS. Volcano plot of ORF9B interactome (n=3) identified by (D)APPLE-MS (138 significant hits) and (E) AP-MS (135 significant hits). In (D) and (E), Differential proteins were identified by two-sided moderated t-tests in Perseus (v2.0). Missing values were imputed using Perseus’ default settings (normal distribution, width=0.3, downshift=1.8). Significant interactors (blue) were defined by FDR-adjusted p < 0.05 and log_2_FC ≥ 1.5, while reported interactors in BioGRID are shown in red. Gray points indicate non-significant proteins. (F)Method comparison. Venn diagram of protein-protein interactions identified by AP-MS (blue), APPLE-MS (green), and literature-curated ORF9B interactors from BioGRID (purple). Database interactions were filtered for ≥2 supporting publications (BioGRID v4.4.244). Circle areas are proportional to total identifications. (G)Schematic of ORF9B complexes with primary (light green) and secondary (dark green) of each method (BioGRID v4.4.244).

To specifically target interactors, we engineered a C-terminally Twin-Strep-tagged ORF9B construct (ORF9B–Twin-Strep-tag) for comprehensive interaction profiling. Immunofluorescence microscopy showed strong mitochondrial localization of ORF9B–Twin-Strep-tag, with significant colocalization observed for the mitochondrial import receptor TOMM70 (also known as TOM70) (Figure 2B). This observation aligns with previous studies identifying TOMM70 as a canonical binding partner of ORF9B^16^. Next, we co-expressed ORF9B–Twin-Strep-tag with Myc-tagged TOMM70 in HEK293T cells and performed an APPLE-MS assay. As expected, the ORF9B–TOMM70 interaction mediated efficient SA-Pup^E^ binding to Myc–TOMM70, confirming the proximity-dependent labeling capability of our APPLE-MS platform (Figure 2C).

We then applied APPLE-MS to systematically characterize the ORF9B interactome, identifying 138 high-confidence interactors (Figure 2D). To benchmark performance, we conducted parallel analyses using conventional AP-MS with Twin-Strep-tag enrichment via streptactin beads (Figure 2E). Comparative analysis (Table S1) revealed three key advantages of APPLE-MS. First, it achieved a 4.07-fold improvement in the number of identified interactors relative to the total number of detected proteins, and it identified twice as many literature-curated interactors from the BioGRID database compared to conventional AP-MS (Figure 2F and S1). Second, whereas conventional AP-MS detected an average of 7,353 proteins per group, our approach confidently identified 4,196 unique proteins per group,with an increased proportion of high-confidence interactors, demonstrating more effective enrichment. Third, APPLE-MS significantly enhanced the signal-to-noise ratio in interaction detection (Table S2). Here, “noise” was defined as data points meeting all of the following criteria: (1) p > 0.05, (2) detection in ≥2 blank controls (untransfected lysates), (3) high inter-replicate variability in both experimental and control groups (CV > 30%) and (4) Absence from traditional MS contamination protein list (CRAPome^17^). This side-by-side comparison enabled a direct assessment of APPLE-MS’s distinct advantages in capturing weak interactions, underscoring its capacity to detect both canonical and novel interactions with high specificity.

We further evaluated each method’s ability to characterize protein complexes associated with ORF9B. Both AP-MS and APPLE-MS identified not only ORF9B’s direct interactors but also secondary complex components, with APPLE-MS providing superior coverage of indirect associations reported in BioGRID (Figure 2G). This improved coverage highlights APPLE-MS’s enhanced ability to map extended protein

### APPLE-MS to identify the dynamic interactome of ORF9B

To demonstrate the applicability of APPLE-MS for capturing dynamic protein-protein interactions (PPIs), we employed this approach to systematically characterize the interactome of SARS-CoV-2 ORF9B. By leveraging the superior sensitivity of APPLE-MS, we aimed to explore how ORF9B interacts with host proteins over time and in response to viral immune activation. To achieve this, we used Poly(I:C), a synthetic analog of double-stranded RNA, which activates pattern recognition receptors, notably RIG-I and MDA5, to trigger potent intracellular antiviral immune responses^18^. As a well-established mimic of viral RNA, Poly(I:C) effectively recapitulates key antiviral defense mechanisms, making it a crucial tool for studying host responses to RNA viruses. Leveraging the superior sensitivity of APPLE-MS, we systematically characterized the dynamic interactome of SARS-CoV-2 ORF9B during a 24-hour Poly(I:C) stimulation time course in HEK293T cells (Figure 3A). This experimental design was guided by the observed temporal upregulation of RIG-I expression (Figure 3B), which served as a marker of successful immune activation.

**Figure 3.**
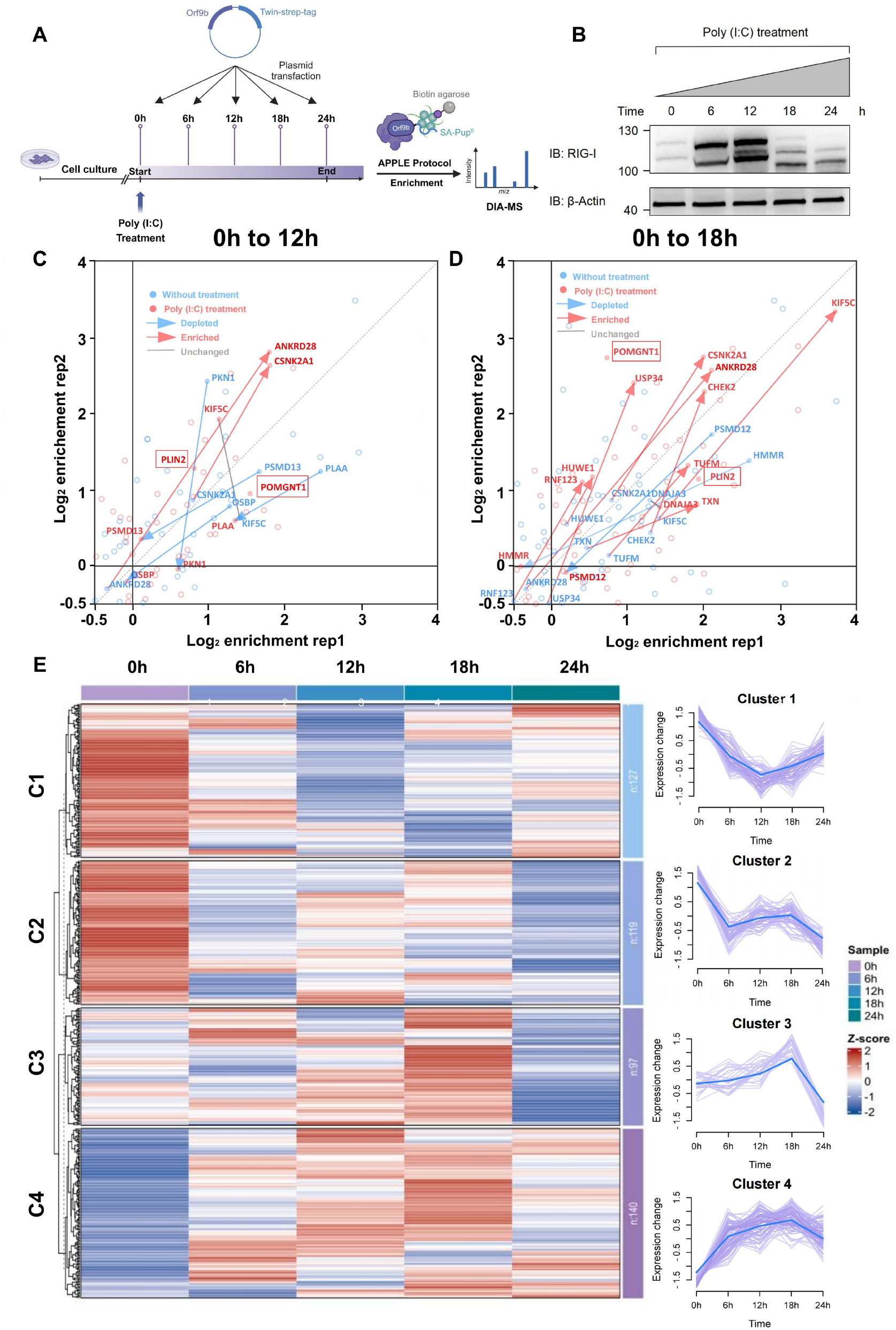
APPLE-MS to identify the dynamic interactome of ORF9B. (A)Workflow of time-course interactome proteomic analysis at five poly(I:C) stimulation time points (0, 6, 12, 18, 24h), with three independent biological replicates collected for each time point. (B)Western blot analysis showing poly(I:C)-induced RIG-I expression dynamics. (C-D) Dynamic changes in the ORF9B interactome following poly(I:C) stimulation at 12h (C) and 18h (D) relative to baseline (0h). In (C) and (D), Scatter plot analysis comparing log_2_ (fold-change) enrichment of ORF9B interactors between untreated (0 h) and poly(I:C)-treated (12h or 18h, 1 μg/mL) conditions demonstrates stimulus-dependent interaction changes, with each point representing protein enrichment in duplicate experiments (x/y-axes: replicate 1/replicate 2 log_2_FC values). Red points represent poly(I:C)-treated samples (n=3 biological replicates), while blue points denote untreated controls. Red arrows highlight proteins with increased association, whereas blue arrows mark those with decreased binding (log_2_FC ≤ -1.5, FDR < 0.05). Gray lines indicate proteins showing no significant change (|log_2_FC| < 1). Statistical thresholds were established by two-tailed moderated t-test (Benjamini-Hochberg adjustment), with dashed diagonal lines marking x=y reference. (E) Temporal clustering of the ORF9B interactome during poly(I:C) transfection. Hierarchical clustering (Left pannel) of log_2_FC values for known (BioGRID) and novel interactors identified in this study, with proteins grouped by mfuzz time-course clustering. The optimal number of clusters (k = 4, m=1.25) was determined by (1) minimum centroid distance (> 0.8) to ensure distinct expression trajectories (2) Knee-point in the SSE curve (Fig. S4), where additional clusters no longer significantly reduced variance. (Right panel) Mfuzz time-course clustering of interaction trajectories, with thick blue lines representing cluster centroids. Each line represents an individual protein.

We identified 104 interactors of ORF9B that were consistently detected across both initial screening and time-course experiments. Analysis of the mass spectrometry data (Figure S2; Table S3) revealed significant temporal shifts in the ORF9B interactome under antiviral conditions. Notably, we observed enhanced interactions between ORF9B and critical components of the RIG-I/DDX58-MAVS signaling cascade, such as the E3 ubiquitin ligase RNF123 and mitochondrial translation elongation factor TUFM. We also identified novel associations with key regulators of apoptosis and autophagy pathways, including casein kinase CSNK2A1 and DNA damage checkpoint kinase CHEK2. Additionally, ORF9B interacted with metabolic regulators, such as lipid droplet protein PLIN2 and glycosyltransferase POMGNT1 (Figure 3C and 3D). In contrast, ORF9B showed reduced binding affinity for core proteasomal subunits PSMD12/13 and the ubiquitin-like modifier PLAA, suggesting an active evasion of host protein quality control systems. These findings highlight ORF9B as a multifunctional virulence hub, capable of modulating host immunity through ubiquitination and phosphorylation networks, reprogramming cellular metabolism, evading proteasomal degradation, and promoting viral replication.

To comprehensively characterize the temporal dynamics of ORF9B interactions, we employed Mfuzz^19^, a soft clustering algorithm for time-series data, to categorize ORF9B interactors into four distinct clusters based on their centroid distances (Figure 3E; Figure S3). This analysis revealed complex biphasic and compensatory interaction patterns during infection. Functional analysis showed that these clusters were predominantly associated with mitochondrial processes, with most identified interactors belonging to clusters 2 and 3. Cluster 2 exhibited a depletion phase (0–6 h), followed by stabilization, and was enriched for TCA cycle enzymes, suggesting that ORF9B may initially target mitochondrial oxidative metabolism to redirect carbon flux toward viral biosynthetic needs. In contrast, Cluster 3 displayed progressive recruitment before abrupt depletion at 24 h, with significant enrichment for oxidative phosphorylation components and metabolic precursor-generating enzymes, indicating that ORF9B exploits energy production for viral particle assembly before inducing late-stage host shutoff. These findings suggest that ORF9B hijacks mitochondrial functions in a temporally coordinated manner. In the early phase of infection, ORF9B disrupts the TCA cycle to accumulate intermediates for viral replication. As infection progresses (6–18 h), ORF9B targets oxidative phosphorylation complexes, particularly components of the electron transport chain, to redirect host energy production toward viral needs. In the late infection phase (>18 h), ORF9B orchestrates a host metabolic shutdown, potentially via inhibition of mitochondrial import machinery or activation of proteolytic pathways. This shutdown may serve dual purposes: suppressing antiviral signaling pathways while facilitating viral particle release. Additionally, we observed that ORF9B modulates mitochondrial gene expression (Cluster 1).

In summary, our study establishes the first time-resolved interactome of ORF9B, uncovering novel binding partners and demonstrating its stage-dependent regulation of host mitochondrial processes during Poly(I:C)-induced antiviral responses. These results provide crucial insights into how SARS-CoV-2 reprograms host metabolism through ORF9B’s phase-specific protein interactions, identifying potential targets for interrupting viral replication cycles.

### APPLE-MS for studying endogenous protein–protein interactions

In endogenous PPI studies, using small affinity tags such as Twin-Strep (∼2–3 kDa) has notable advantages compared to employing larger proximity labeling enzymes (∼20–50 kDa). Their compact size simplifies genetic insertion and minimizes potential interference with the protein’s native function. These features make methods like APPLE-MS particularly suitable for mapping PPIs under near-physiological conditions. In endogenous PPI studies, small affinity tags such as Twin-Strep (∼2–3 kDa) offer notable advantages over larger proximity labeling enzymes (∼20–50 kDa). Their compact size simplifies genetic insertion and minimizes potential interference with the protein’s native function. These features make methods like APPLE-MS particularly suitable for mapping PPIs under near-physiological conditions. PIN1, a phosphorylation-specific prolyl isomerase that catalyzes the cis-trans isomerization of phosphorylated Ser/Thr-Pro motifs, serves as a critical regulator of cell cycle progression, transcriptional control, and neurodegenerative pathways^20^. To comprehensively characterize the PIN1 interactome under near-physiological expression conditions, we used CRISPR-Cas9 to insert a Twin-Strep-tag at the native PIN1 locus, facilitating extensive protein-protein interaction (PPI) mapping via APPLE-MS at endogenous expression levels (Figure 4A-C). Comparative analyses clearly demonstrated the superior performance of APPLE-MS over conventional affinity purification mass spectrometry (AP-MS) (Figure 4D and 4E), revealing significantly more known PIN1 interactors, including low-abundance regulators such as ITGA4, with improved signal-to-noise ratios and greater enrichment (Figure 4F and 4G; Table S4). These findings underscore APPLE-MS’s enhanced sensitivity for delineating endogenous protein networks under native physiological conditions.

**Figure 4.**
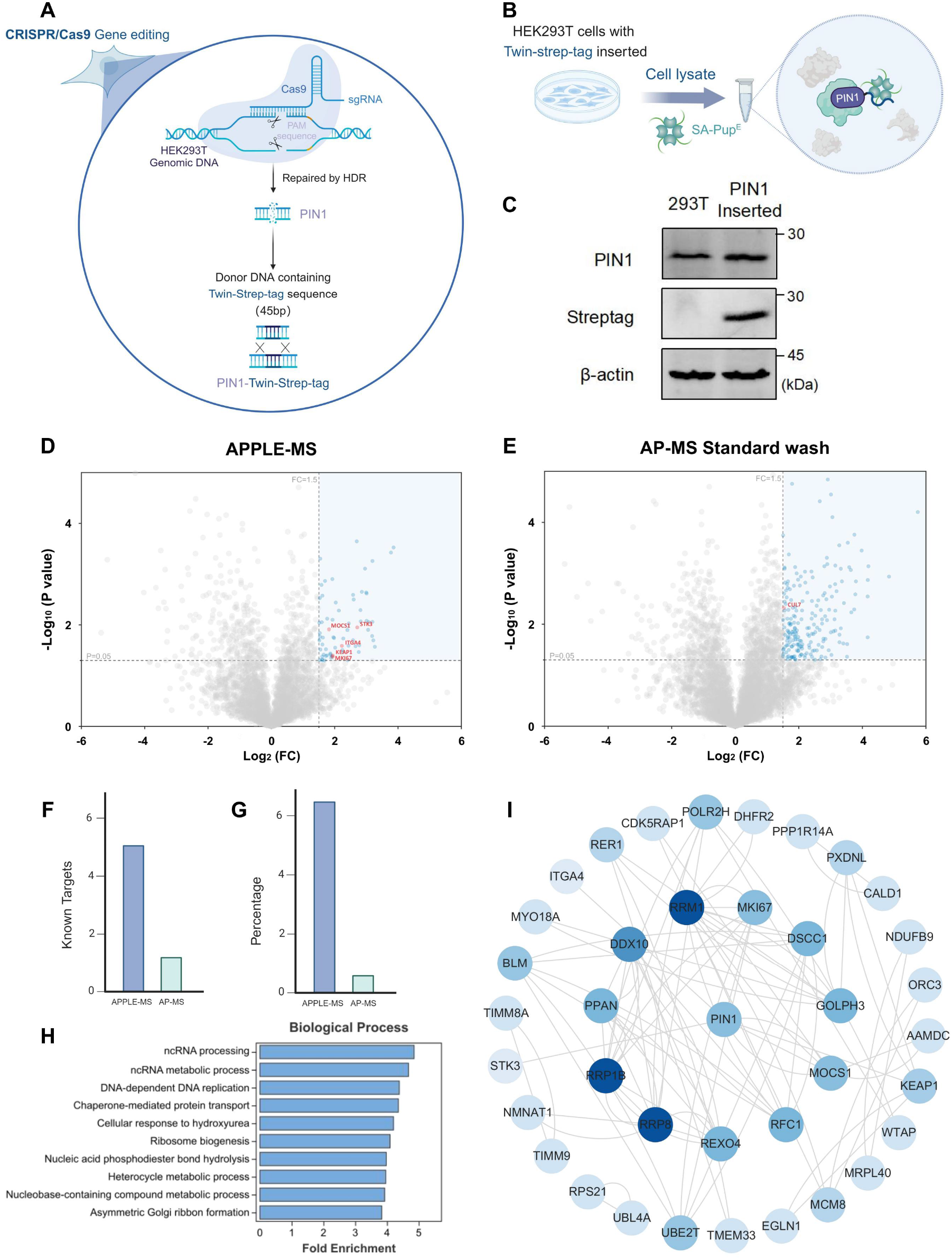
APPLE-MS to study endogenous proteins. (A)Genome engineering strategy. HDR-mediated C-terminal Twin-Strep-tag insertion at the PIN1 locus. (B)Endogenous PIN1 Interaction Profiling by APPLE-MS. (C)Tag validation. Western blot confirming Twin-Strep-tag insertion. (D-E) PIN1 interactome profiling. Volcano plot of PIN1 interactome (n=3) identified by (D) APPLE-MS (78 significant hits) and (E) AP-MS (200 significant hits). In (D) and (E), Differential proteins were identified by two-sided moderated t-tests in Perseus (v2.0). Missing values were imputed using Perseus’ default settings (normal distribution, width=0.3, downshift=1.8). Significant interactors (blue) were defined by FDR-adjusted p < 0.05 and log_2_FC ≥ 1.5, while reported interactors in BioGRID are shown in red. Gray points indicate non-significant proteins. (F-G) Method comparison. (F) Absolute counts and (G) detection rates of known PIN1 interactors. (H)Functional enrichment. Bar plot displays significantly enriched biological processes (FDR < 0.05, hypergeometric test) among PIN1 interactors identified by APPLE-MS (n = 3). The length of each bar corresponds to the -log_10_ (FDR) value, representing the statistical significance of enrichment. (I)PIN1 interaction network topology. Cytoscape visualization of PIN1-associated proteins (nodes) and their interactions (edges). Node color intensity scale with degree (range: 1–25), highlighting proteins for subsequent verification.

Gene Ontology (GO) enrichment analysis (Figure 4H and S4) of the APPLE-MS-derived PIN1 interactome identified two functionally significant pathways beyond its canonical roles: ncRNA processing and DNA replication machinery. Enrichment of DNA replication components (e.g., RFC1, MCM complex) aligns with prior reports indicating PIN1’s S-phase-specific nuclear localization^21^ and its role in replication fork stabilization^22^. These results reinforce PIN1’s function as a guardian of genomic stability, where its prolyl-isomerase activity modulates replication stress responses by facilitating fork protection complexes, coordinating checkpoint signaling during replication stress, and sustaining replisome component activity under genomic insult. Additionally, identifying MNAT highlights PIN1’s broader involvement in cell cycle regulation, notably at the G1/S transition (Figure 4I). These discoveries position PIN1’s isomerase activity as a spatiotemporal regulator that simultaneously ensures genomic fidelity and fine-tunes gene expression networks, particularly significant in oncogenic and neurodegenerative contexts where both processes are commonly dysregulated.

### APPLE-MS to identify membrane interactors

To evaluate our method’s capability for *in situ* identification of cell-surface PPIs, we employed the APPLE-MS assay to profile the interactome of glucagon-like peptide-1 (GLP-1), a hormone critically relevant to diabetes and obesity management (Figure 5A). Given GLP-1’s pleiotropic effects beyond glycemic control^23^, a detailed characterization of its receptor complexes could identify novel therapeutic targets and mechanisms underlying its extra-pancreatic actions. For validation, we established a model system by transiently transfecting HEK293T cells with a GLP-1 receptor (GLP-1R) expression plasmid. Notably, all proximity labeling reactions were conducted directly on adherent cells in culture dishes, maintaining native membrane contexts, physiological receptor conformation, and endogenous interaction kinetics for PPI capture. Western blot analysis confirmed SA-Pup^E^ conjugation to GLP1-R (Figure 5B), and immunofluorescence assays validated GLP-1-Twin-Strep-tag/GLP-1R complexes at the plasma membrane (Figure 5C), confirming efficient labeling capability and interaction specificity. Subsequently, mass spectrometry effectively identified membrane interactors, including GLP1-R in overexpressing cells, affirming APPLE-MS’s sensitivity for membrane proteins (Figure 5D; Table S5). Our analysis also detected multiple GLP-1-associated proteins from the BioGRID database. We propose two plausible explanations for these interactions: either these proteins represent endogenous GLP-1-binding partners unaffected by GLP-1R transfection, or they indirectly associate with GLP-1 through receptor-mediated interactions, possibly forming ternary complexes or participating in GLP-1R-dependent signaling networks, positioning GLP-1R as a central regulator of multiprotein assemblies.

**Figure 5.**
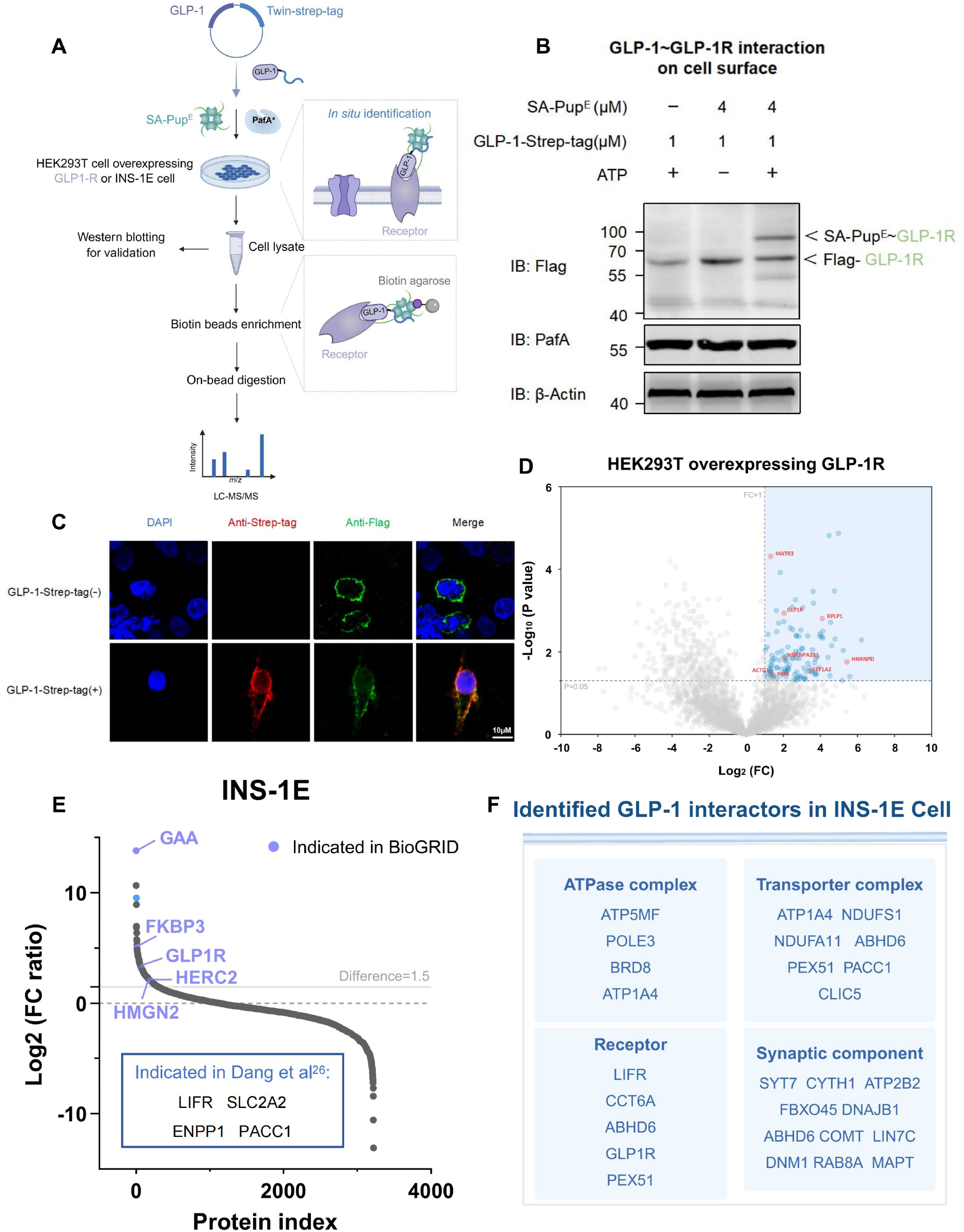
APPLE-MS to identify membrane interactors. (A)Schematic design of APPLE-MS for identifying cell-surface protein interactors. The labeling process is performed *in situ* by removing the culture medium from the dish. Interactions between the Twin-Strep-tagged bait and prey protein are then detected via gel electrophoresis or mass spectrometry. (B)Western blot analysis of *in situ* GLP-1-GLP-1R interaction in HEK293T cells using the APPLE assay. (C)Immunofluorescence imaging demonstrating cell surface colocalization (yellow) of GLP-1R (green) with GLP-1-Twin-Strep-tag (red) in HEK293 cells expressing GLP-1R. Anti-strep-tag were used to stain GLP-1-Twin-Strep-tag. Nuclei were counterstained with DAPI (blue). Representative images from three independent experiments are shown. Scale bars, 10 µm. (D)GLP-1 interactome profiling by APPLE-MS. Volcano plot of GLP-1 interactome (96 significant hits). Differential proteins were identified by two-sided moderated t-tests in Perseus (v2.0). Missing values were imputed using Perseus’ default settings (normal distribution, width=0.3, downshift=1.8). Significant interactors (blue) were defined by FDR-adjusted p < 0.05 and log_2_FC ≥ 1, while reported interactors in BioGRID are shown in red. Gray points indicate non-significant proteins(n=3). (E)Rank plot analysis of log_2_FC ratio values of proteins in two biological replicates. Reported interactors in BioGRID are highlighted in red. Indicated interactors in Dang et al. are highlighted in blue. (F)Functional categorization. Representative GLP-1R interactors from INS-1E cells grouped by biological function.

Next, we applied APPLE-MS to INS-1E cells, a well-characterized rodent pancreatic β-cell line expressing endogenous GLP-1R and native membrane polarity. Our analysis identified 301 putative interactors, including five previously reported GLP-1R-associated proteins from BioGRID and four indicated in Dang et al^28^. (Figure 5E; Table S6). Notably, our method uncovered unexpected co-receptors and transporter complexes (Figure 5F), demonstrating its potential for revealing novel GPCR signaling network components. The interactome analysis highlighted a complex GLP-1 signaling network featuring functionally significant clusters. Key ATPase components (ATO5MF, BRD8, POLE3 and ATP1A4) and transporter complexes (ABHD6, CLIC5, NDUFS1, NDUFA11, etc) potentially coordinate ion homeostasis and energy metabolism, with ATP1A4 emerging as a possible molecular bridge between membrane potential regulation and GLP-1-dependent insulin secretion. The detection of receptor-associated proteins such as DNM1 and GLP-1R implies sophisticated regulation of receptor endocytosis and cAMP/PKA signaling kinetics, consistent with GLP-1’s established role in glucose-stimulated insulin secretion^24^.

These findings significantly enhance our mechanistic understanding of GLP-1 by revealing its involvement in membrane dynamics and neuro-metabolic interfaces. The APPLE-MS approach, leveraging proximity labeling, robustly captured context-specific interactors, providing an effective platform for future studies of GLP-1’s tissue-specific roles. Notably, while our current work focused on steady-state interactome, the APPLE-MS platform enables future time-resolved studies of GLP-1R or other membrane receptors through controlled addition of Twin-Strep-tagged ligands.This methodological advancement facilitates systematic exploration of ligand-dependent receptor interactome dynamics in native cellular environments.

## DISCUSSION

In this study, we introduced APPLE-MS and demonstrated its effectiveness in revealing PPIs that are elusive to traditional methodologies. **APPLE-MS synergizes the complementary strengths of AP-MS and Proximity Labeling**, enabling detection of a more complete interactome for a given bait protein. To systematically evaluate its advantages, we compared APPLE-MS with traditional affinity purification mass spectrometry (AP-MS) and proximity labeling techniques (BioID/TurboID/APEX) (Table 1). By combining the high specificity of Twin-Strep-tag enrichment with the proximity-dependent labeling capability of PafA, APPLE-MS effectively captures weak, transient, and membrane-associated interactions in their native cellular context. Our study demonstrates that APPLE-MS provides critical improvements over existing methods, including increased sensitivity for transient interactions, superior signal-to-noise ratios, and improved interactome coverage. The compatibility of APPLE-MS with physiological protein expression levels, demonstrated by CRISPR tagging of PIN1, further minimizes potential experimental artifacts. This work builds upon our previous work^25^ development of SPIDER(Specific Pupylation as IDEntity Reporter), which utilized PafA-mediated PupE labeling coupled with the streptavidin-biotin system to efficiently capture protein interactions with diverse biomolecules (e.g., nucleic acids, small molecules), including m6A-binding proteins and SARS-CoV-2 membrane receptors. While SPIDER excelled in *in vitro* applications, its reliance on biotinylation limited its utility for in vivo interactome profiling. By replacing biotin with the Twin-Strep-tag, APPLE-MS overcomes this constraint, enabling in situ labeling of endogenous protein complexes while retaining the covalent linkage advantages of Pup^E^.

**Table 1.**
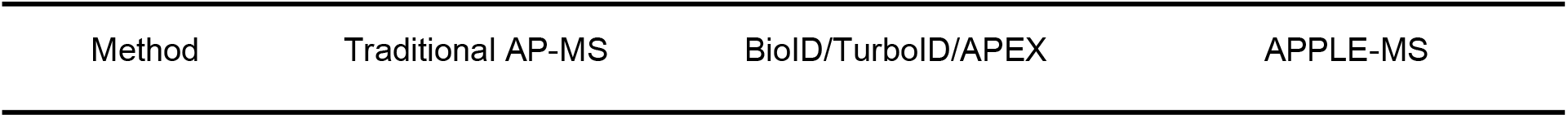
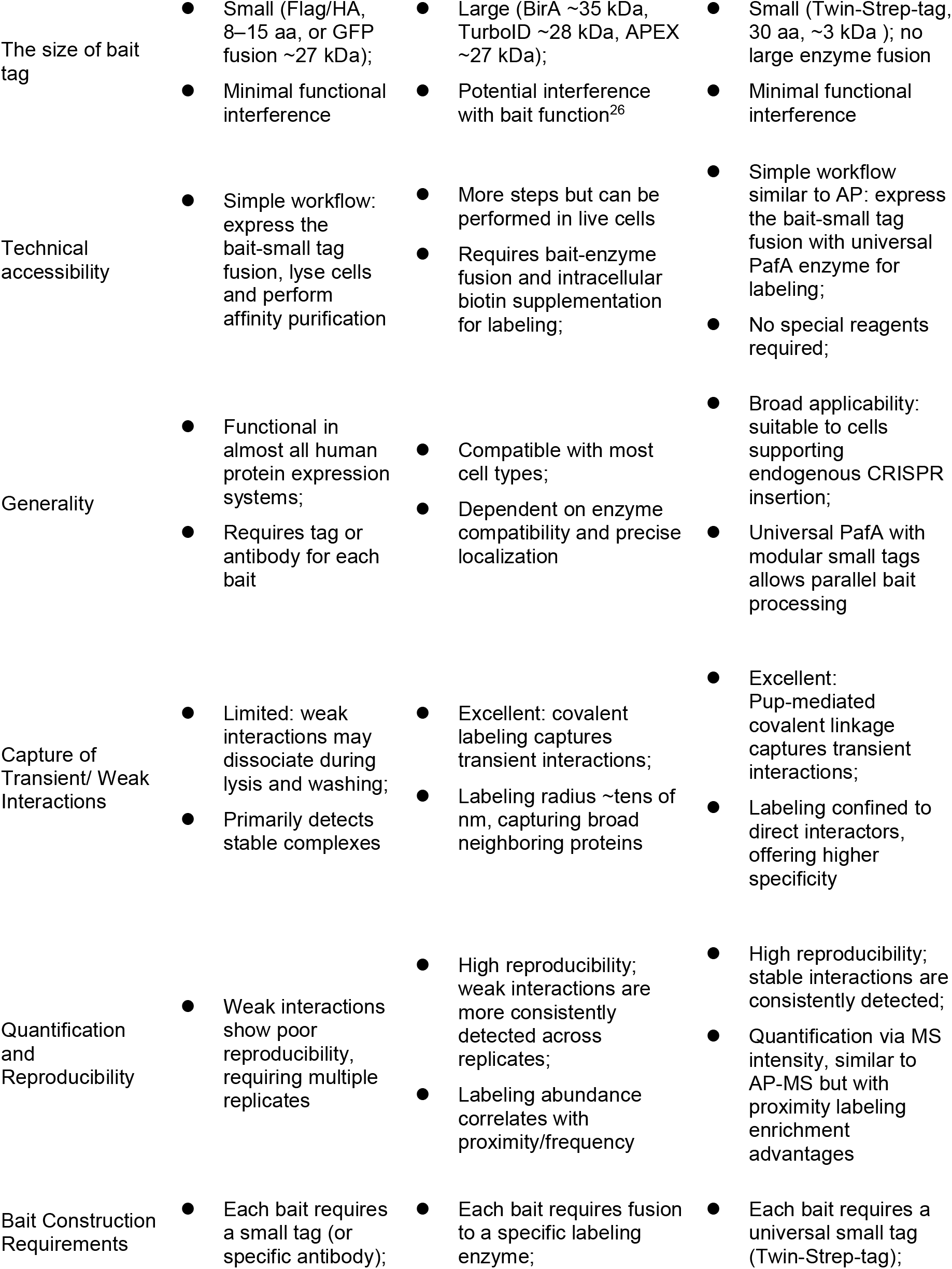
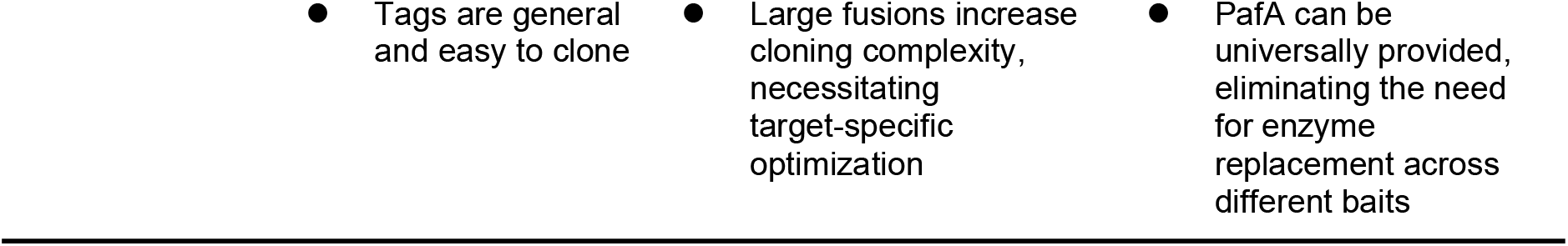
Comparison between traditional AP-MS, BioID/TurboID/APEX and APPLE-MS.

Using the well-characterized SPOP MATH domain-peptide interaction model, we established that APPLE-MS can reliably detect interactions with dissociation constants (Kd) up to 76 μM. This sensitivity significantly surpasses conventional AP-MS, which typically fails to identify interactions weaker than ∼10 μM due to the noncovalent nature that could not endure stringent washing steps^27^. The covalent labeling mechanism of PafA ensures the stable capture of even low-abundance or transient interactors, addressing a key limitation of non-covalent approaches. Direct comparisons with AP-MS further validated APPLE-MS’s enhanced capability, as exemplified by a 4.07-fold increase in high-confidence interactors for SARS-CoV-2 ORF9B with reduced non-specific background. Additionally, APPLE-MS exhibited superior performance in detecting secondary complex components, such as those involved in RNA processing, thus providing deeper insights into complex protein networks. Moreover, APPLE-MS enables the *in situ* labeling of cell-surface proteins, such as the GLP-1 receptor (GLP-1R), without the need for cell lysis, preserving physiologically relevant membrane contexts and interactions. This is particularly advantageous for studying membrane receptors, where interactions are often dependent on native structural integrity.

Our interactome analyses provided new biological insights across diverse targets. For SARS-CoV-2 ORF9B, APPLE-MS revealed multifaceted roles in viral pathogenesis, including mitochondrial dysfunction, immune evasion, and RNA processing manipulation. Dynamic interactions with key metabolic enzymes and immune regulators, such as RNF123 and TUFM, highlighted ORF9B’s complex strategy in reshaping host cellular processes during infection. Similarly, applying APPLE-MS to endogenous PIN1 uncovered involvement in DNA replication stress responses and non-canonical RNA processing pathways. The identified associations, including the MCM complex and RFC1, underscore PIN1’s broader regulatory roles beyond its canonical prolyl-isomerase function. For GLP-1R, APPLE-MS identified canonical and previously unexplored co-receptors and signaling components, revealing new potential regulatory mechanisms and therapeutic targets for diabetes and obesity.

Despite its strengths, APPLE-MS has certain limitations. Non-specific labeling, although significantly minimized by optimized reaction conditions, can still introduce background noise. Strategies such as SILAC or TMT labeling could further enhance the specificity by enabling stringent comparative analyses. Additionally, CRISPR-based endogenous tagging, despite providing physiological relevance, involves challenges such as extensive clonal validation and potential off-target genomic effects.

Overall, APPLE-MS bridges critical gaps in interactome mapping by integrating the complementary strengths of AP-MS and proximity labeling methods. It is broadly applicable to intracellular, membrane, and organellar interactomes across various biological contexts. Future developments, including multiplexed labeling strategies, adaptations for primary cells and tissues, and integration with structural proteomics methods such as cryo-EM, promise to further enhance the method’s utility and impact.

## RESOURCE AVAILABILITY

### Lead contact

Further information and requests for resources and reagents should be directed to and will be fulfilled by the lead contact, Hewei Jiang.

### Materials availability

All unique plasmids and/or reagents generated in the study are available from the [lead contact](https://www.cell.com/cell-reports/fulltext/S2211-1247(25)00201-3) with a completed materials transfer agreement.

### Data and code availability

- Original western blot images and microscopy data reported in this paper will be shared by the lead contact upon request.
- The mass spectrometry proteomics data have been deposited to the ProteomeXchange Consortium via the PRIDE partner repository with the dataset identifier PXD063145.
- Any additional information required to reanalyze the data reported in this paper is available from the lead contact upon request.

## ACKNOWLEDGMENTS

This work was funded by Lingang Laboratory (Startup Fund) and partially supported by the Fourteenth Five-Year National Key Research and Development Program of China (2023YFC2307200), the Natural Science Foundation of China (No. 92374110 and 32271492), the R&D Program of Guangzhou National Laboratory (No. GZNL2023A01005).

## AUTHOR CONTRIBUTIONS

Conceptualization, H.W.J. and S.C.T.; Methodology/Investigation, S.L., L.X., L.Y., Z.H., L.W. and Y.W.; Writing, S.L..and H.J.; Resources, H.Z and S.G.

## SUPPLEMENTAL INFORMATION

**Document S1. Figures S1–S4**

**Table S1. Comparative proteomic profiling of ORF9B interactors in 293T cells using APPLE-MS and AP-MS**.

**Table S2. Noise protein lists identified by APPLE-MS and AP-MS**.

**Table S3. Temporal proteomic analysis of ORF9B interactors under poly(I:C) stimulation.**

**Table S4. Comparative interactome mapping of PIN1 in 293T cells via APPLE-MS and AP-MS.**

**Table S5. GLP-1 interactome profiling in 293T cells overexpressing GLP-1R**.

**Table S6. Cell-specific interactome of GLP-1 in INS-1E cells**.

## EXPERIMENTAL MODEL AND STUDY PARTICIPANT DETAILS

### Cell culture

APPLE-MS was tested on HEK 293T and INS-1E cells. HEK293T cells were cultured in DMEM (Thermo Fisher Scientific) and INS-1E cells were cultured in RPMI 1640 (Thermo Fisher Scientific). The culture mediums were supplemented with 10% (v/v) heat-inactivated fetal bovine serum (FBS) and 1% Penicillin-Streptomycin. All cell lines were cultured in 37°C and 5% CO_2_. Cell passaging was conducted at 2-4day intervals. No mycoplasma contamination was detected for all cell lines.

## METHOD DETAILS

### Plasmids construction and cloning

Protein sequences were downloaded from Addgene. The corresponding DNA sequences were codon-optimized and synthesized by Tsingke Biotech (Shanghai, China), and the synthesized genes were cloned into pET28a or pET23a for prokaryotic expression or pcDNA3.1 for eukaryotic expression. All plasmids were constructed using homologous recombination. Target genes were amplified by PCR (Phanta SE Super-Fidelity DNA Polymerase, Vazyme) with primers containing 15–20 bp overlapping regions. Ligated products were transformed into chemically competent E. coli DH5α or BL21 (Sangon Biotech) plated on LB-agar with appropriate antibiotics, and validated by Sanger sequencing (Tsingke Biotech).

### Protein expression and purification

E. coli BL21, carrying the expression plasmid, was cultured in LB medium at 37°C until the A_600_ reached 0.6∼0.8. Protein expression was induced with 0.5 mmol L−1 isopropyl-β-D-thiogalactoside (IPTG) at 18°C for 16 hr. Cell pellets expressing 6×His-tagged proteins were resuspended in lysis buffer (50 mM Tris-HCl, pH 8.0, 500 mM NaCl, 20 mM imidazole) and lysed using a high-pressure homogenizer (Union-Biotech, China). The lysate was clarified by centrifugation (10,000 × *g*, 20 min, 4°C), and the supernatant was incubated with Ni-NTA Agarose (Qiagen). After binding, the resin was washed with lysis buffer and eluted with an imidazole gradient (300 mM in 50 mM Tris-HCl, pH 8.0, 500 mM NaCl).

### Cell transfection and lysate preparation

For transfection, we employed polyethylenimine (PEI, 1 mg/mL, Polysciences) at a ratio of 2 μL PEI per 1 μg plasmid DNA (e.g., 4 μL PEI for 2 μg plasmid). The plasmid DNA was first diluted in 50 μL of serum- and antibiotic-free DMEM medium and mixed gently by pipetting. In parallel, the appropriate amount of PEI was diluted in another 50 μL of serum- and antibiotic-free DMEM, while the corresponding amount of PEI was separately diluted in another 50 μL of serum-free DMEM. Within 5 minutes of preparation, the plasmid and PEI solutions were combined, thoroughly mixed, and incubated at room temperature for 15 minutes to allow optimal DNA-PEI complex formation. These complexes were then added dropwise to 293T cells that had reached approximately 85% confluency. Following 48 hours of incubation to allow protein expression, transfected cells were processed for either immediate analysis or cell lysis.

For protein extraction, transfected cells were first washed with PBS before lysis with 1 mL of M-PER Reagent (Thermo Fisher Scientific) supplemented with 0.5 mM PMSF per 10 cm culture dish. Complete lysis was achieved by gentle shaking for 20 minutes at room temperature. The resulting lysate was then clarified by centrifugation at 12,000×g for 10 minutes to remove cellular debris. The protein-rich supernatant was carefully aliquoted and stored at -80°C to preserve protein integrity for subsequent downstream applications.

### APPLE-MS assay

For the APPLE experiment validating the interaction between bait peptides and GFP-MATH, Twin-Strep-tag-fused bait peptides 1-3, GFP-MATH, SA-Pup^E^, PafA12KR and ATP were co-incubated at 37°C for 45 min in the reaction buffer (50 mM Tris-HCl, 150 mM NaCl, 20 mM MgCl_2_). The concentrations used were 1 μM for each Twin-Strep-tag-fused bait peptide (1-3), 1 μM for GFP-MATH, 10 μM for SA-Pup^E^, 10 mM for PafA12KR and 5 mM for ATP. Reactions were terminated by adding SDS loading buffer and boiling at 95°C for 10 min. The supernatant was then collected by centrifugation for WB detection.

For the conventional APPLE assay to identify PPIs, 3mg of cell lysate transfected with plasmid expressing the Twin-Strep-tag-fused bait protein was incubated in reaction buffer at 37°C for 45 min. The final concentrations of reagents were 0.25 μM SA-Pup^E^, 0.25 μM PafA12KR and 5 mM ATP, with 0.5 mM PMSF added to prevent protein degradation. For mass spectrometry, reactions were stopped by adding 8M urea. The cell lysate was incubated with continuous rotation at room temperature until complete urea dissolution. 100 μL biotin agarose (Sigma-Aldrich) was added to the supernatant and incubated at 4°C overnight.

For APPLE assay on the cell surface, the culture medium was first removed from the dish, and cells were washed twice with pre-warmed PBS (37°C). A reaction mixture containing 0.2 μM GLP-1-Twin-Strep-tag, 0.2 μM SA-Pup^E^ and reaction buffer supplemented with 0.5mM PMSF was prepared in a 15 mL conical tube and incubated at room temperature for 30 min. This mixture was then added to cells in dish and incubated at 4°C for 1-2 h. Finally, 0.2 μM PafA and 5 mM ATP were added to initiate the enzymatic reaction, which proceeded at 37°C for 40 min. All steps were performed under sterile conditions with PMSF added immediately before use to maintain protease inhibition activity. After reaction completion, cells were lysed using RIPA buffer. After complete cell lysis, 8 M urea was added to the lysate to a final concentration of 8 M. The protein interaction complexes were then isolated using the same affinity capture procedure described above.

### Immunoblotting

Following electrophoretic separation on 7.5-12.5% gradient SDS-polyacrylamide gels, proteins were transferred to methanol-activated PVDF membranes (Merck Millipore) using a wet transfer system (Mini Trans-Blot, Bio-Rad). The transfer buffer (25 mM Tris-HCl, 190 mM glycine, 20% methanol) was maintained at room temperature during 1.5 h transfer at 160 mA constant current. Membranes were then briefly rinsed with TBST and blocked for 2 hour at room temperature with 5% non-fat dry milk (Sangon Biotech) in TBST. All primary and secondary antibodies (detailed in KEY RESOURCES TABLE) were diluted in TBST and applied with three TBST washes between incubations. The membrane was scanned on an imager system (Typhoon).

### Immunofluorescence

After a 48 hours post-transfection, cells were washed twice with pre-warmed PBS and fixed with 4% paraformaldehyde (PFA) for 15-20 min at room temperature (protected from light). After three PBS washes, cells were permeabilized with 0.2% Triton X-100 in PBS for 10 min at room temperature and washed again three times with PBS. Non-specific binding was blocked overnight with 5% BSA in PBS at 4°C. Cells were then sequentially incubated with primary antibodies and corresponding fluorescent secondary antibodies, with thorough PBS washes between each step. Nuclei were counterstained with DAPI-containing antifade mounting medium, and coverslips were sealed with nail polish. Fluorescence images were acquired using a confocal microscope.

### Affinity purification for protein-protein interactions

30 μL STarm Streptactin Beads 4FF (Smart-Lifesciences Biotechnology, SA053005) were added to 3 mg cell lysate transfected with plasmid expressing the Twin-Strep-tag-fused bait protein in binding buffer (50 mM Tris-HCl, 150 mM NaCl, 20 mM MgCl_2_, pH 7.4) and incubated overnight at 4°C with rotation. After incubation, beads were washed twice with PBS to remove nonspecifically bound proteins. The washed beads were then directly processed for mass spectrometry.

### Poly (I:C) treatment

For Poly I:C transfection, Lipofectamine™ 3000 Transfection Reagent (Thermo Fisher Scientific, L3000001) was employed according to the manufacturer’s protocol with modifications. Briefly, for each 10 cm culture dish, 24 μg of Poly I:C (Medlife, PC11566) was diluted in 500 μL of serum- and antibiotic-free DMEM medium and mixed gently by pipetting, followed by addition of 36 μL P3000™ Reagent. In parallel, 48 μL of Lipofectamine 3000 reagent was diluted in another 500 μL of serum-free DMEM. The two solutions were combined within 5 min, mixed thoroughly, and incubated at room temperature for 15 min to allow complex formation. The resulting transfection complexes were then added dropwise to 293T cells that had reached approximately 85% confluency.

### Coimmunoprecipitation

Protein-protein interactions were investigated through co-immunoprecipitation assays using HEK293 cells transiently expressing ORF9B-Twin-Strep-tag and HA-tagged candidate interactors. Following cell lysis, cleared supernatants containing 2 mg total protein were subjected to affinity purification using STarm Streptactin Beads 4FF (Smart-Lifesciences Biotechnology, SA053005) overnight at 4°C. After two washes with PBS, bound proteins were eluted by heat denaturation in SDS loading buffer (95°C, 10 min) and analyzed by immunoblotting with anti-strep-tag and anti-HA antibodies.

### Mass spectrometry

Biotin-agarose enriched proteins were subjected to reduction with 10 mM DTT (37°C, 1 h) followed by alkylation using 25 mM iodoacetamide (dark incubation, 20 min). On-bead tryptic digestion was then performed at 37°C overnight using a 1:30 (w/w) enzyme-to-protein ratio. After digestion, the beads were washed twice with 200 μL of 50 mM ammonium bicarbonate (NH_4_HCO_3_). The combined supernatants containing tryptic peptides were desalted using Empore StageTips6094 SDB-RPS (MGAM Institute, YW-SDB-RPS-96) following the manufacturer’s protocol.

The peptides were re-dissolved in solvent A (A: 0.1% formic acid in water) and analyzed by Orbitrap Exploris 480 coupled to an EASY-nanoLC 1200 system (Thermo Fisher Scientific, MA, USA). 3 μL peptide sample was loaded onto a 25 cm analytical column (75 μm inner diameter, 1.9 μm resin (Dr Maisch)) and separated with 60 min-gradient starting at 4% buffer B (80% ACN with 0.1% FA) followed by a stepwise increase to 50% in 53 min 40 s, 95% in 40 s and stayed there for 5 min 40 s. The column flow rate was maintained at 300 nL/min with the column temperature of 40°C. The electrospray voltage was set to 2 kV. The mass spectrometer was run under data independent acquisition (DIA) mode, and automatically switched between MS and MS/MS mode. The survey of full scan MS spectra (m/z :350-1200) was acquired in the Orbitrap with 120,000 resolutions. The Normalized automatic gain control (AGC) target of 300% and the maximum injection time of 50 ms. Then the precursor ions were selected into collision cell for fragmentation by higher-energy collision dissociation (HCD), the normalized collection energy was: 25%, 30%, 35%. The MS/MS resolution was set at 30,000, the Normalized automatic gain control (AGC) target of 200%, the maximum injection time of 50 ms.

### Proteomics data analysis

Tandem mass spectra were processed by PEAKS Studio version 12 (Bioinformatics Solutions Inc., Waterloo, Canada). The databases were uniprot-homo_sapiens (version 2024, 20608 entries) and uniport-Rattus_norvegicus(version 2024, 22897 entries) which download from uniprot. Trypsin and Lys C was set as the digestion enzyme and semi-Specific was specified as the digest type. PEAKS DB were searched with a fragment ion mass tolerance of 0.02 Da and a parent ion tolerance of 10 ppm. The max missed cleavages was 2. The proteins with 1%FDR and containing at least 1 unique peptide were filtered.

In our experiments, the contaminant standards were proteins that appeared in >50%-70% of AP-MS experiments in the CRAPome database (version 2.0). Putative ORF9B interactors were clustered based on their kinetic profiles using Mfuzz time-course analysis. Protein interaction data were integrated from two public databases: BioGRID (version 4.4.244) for curated interactions and STRING (version 12.0) with a medium-confidence threshold (0.4). These datasets were used to generate a comprehensive cell membrane interactome map through Cytoscape platform (version 3.10.1).

### Generation of Endogenous Tagging of PIN1 Knock-In Cell Line

Addition of PIN1 gene endogenously labelled tag sequence within the genome of HEK293T cells was performed using CRISPR/Cas9 gene editing. Guide RNAs(gRNAs) were designed and cloned into a Cas9-mCherry expression vector to target the insertion site. For the donor construct, 800 base pair homology arms that span left and right of the start codon of PIN1 tagging were generated by Gibson assembly. HEK293T cells were plated in 6-well plates to reach 70% confluence on the next day for transfection. The gRNA plasmid (1.5μg) and donor construct (1.5μg) were introduced into HEK293T cells per well. After 48h of transfection, mCherry positive cells were sorted into 96 well plates to generate single cell clones. To confirm HDR knock-in, monoclonal cell lines were genotyped via the gDNA extraction and PCR amplification. For wild type cells, the band corresponds to a 1010bp product, whereas the insertion of Tag results in a 1142bp product.

### Structure analysis

The protein structures were retrieved from the Protein Data Bank (PDB; https://www.rcsb.org/). Structural examinations, including surface representation and electrostatic potential analysis, were executed using PyMOL Molecular Graphics System (Version 4.6.0, Schrödinger LLC) with default rendering settings.

## QUANTIFICATION AND STATISTICAL ANALYSIS

All statistical analyses were performed using GraphPad Prism (version 10.2.0). The number of replicates (n) of each experiment can be found in the respective figure caption, and n represents the number of independent shake flask cultures. Quantitative data were processed using ImageJ and MaxQuant, with normalization to appropriate controls. Multiple testing corrections (Benjamini-Hochberg FDR) were applied where appropriate, with significance thresholds set at P < 0.05. Significant m/z features were defined with a 1.5-fold cut-off and a *p*-value<0.05 for a two-sample t-test.

