## Supplemental Figure 1-4 for "APPLE-MS: A affinity purification-mass spectrometry method assisted by PafA-mediated proximity labeling"

**Figure S1**

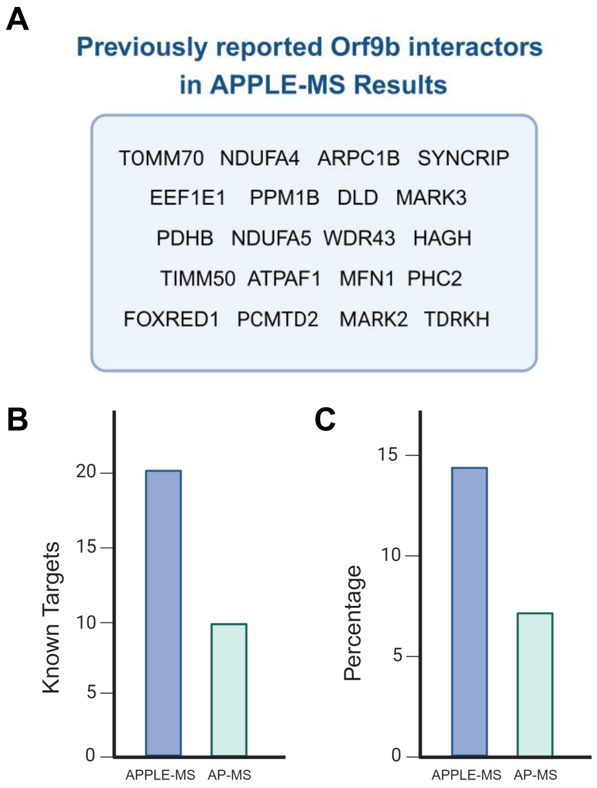


**Figure S1 Comparative analysis of ORF9B interactome detection by APPLE-MS versus AP-MS**

**(A)** Reported ORF9B interactors from BioGRID (v4.4.244) detected by APPLE-MS

**(B)** Quantitative assessment of reported ORF9B interactors. Bar plot illustrates the absolute number of benchmark ORF9B interactors identified by APPLE-MS (blue) versus conventional AP-MS (green).

**(C)** Detection efficiency for reported ORF9B interactors. Bar plot shows the percentage of known interactors among total identified putative binding partners for each method.

### Figure S2

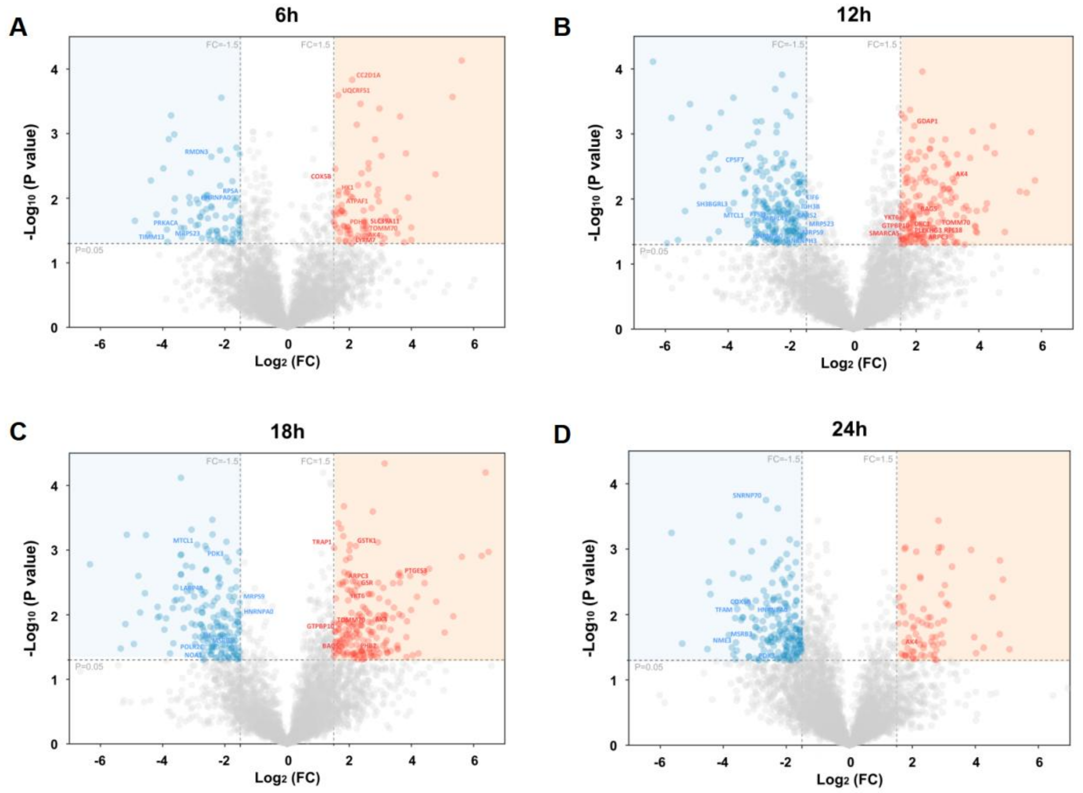


**Figure S2 Temporal Profiling of the ORF9B Interactome Following Poly(I:C) Transfection**

**(A-D)** Time-resolved analysis of ORF9B-protein interactions after poly(I:C) transfection. Volcano plots display significantly enriched interactors ($p < 0.05$, fold-change > 2) at each time point (6, 12, 24, and 48 h; n = 3 biological replicates). Literature-curated ORF9B interactors from BioGRID (v4.4.244) are highlighted in red. Dashed lines indicate statistical thresholds (horizontal: $p = 0.05$; vertical: 1.5-fold change).

**Figure S3**

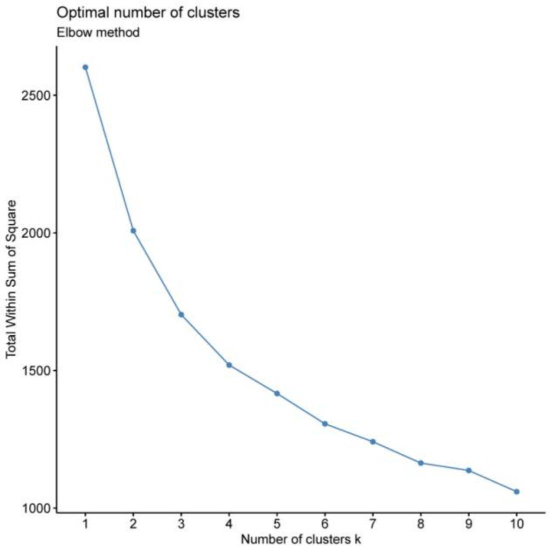


**Figure S3 Determination of Optimal Cluster Number in Mfuzz Analysis**

Line plot of SSR values versus cluster number (k = 1–10) computed from Mfuzz soft clustering of ORF9B interactome dynamics (0–24 h poly(I:C)). The elbow point at k = 4 was selected as the optimal cluster number, balancing model fit and biological interpretability.

**Figure S4**

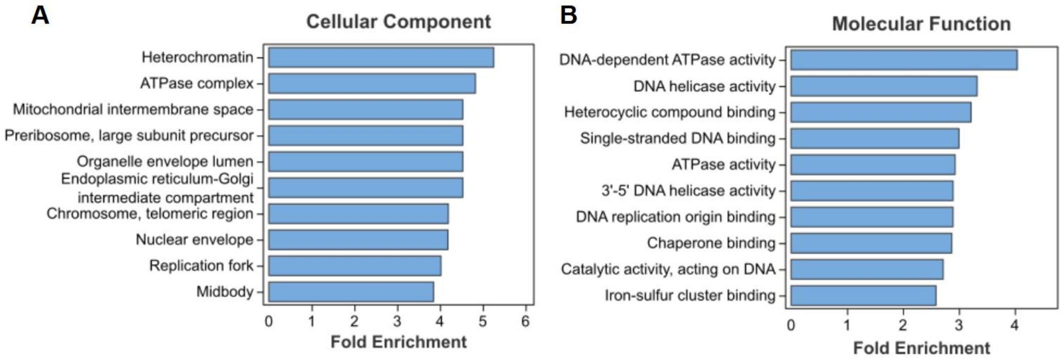


**Figure S4 Cellular component and molecular function of PIN1 putative interactors.**

(A) Subcellular localization of ORF9B interactome. Bar plot shows significantly enriched cellular components (FDR<0.05, hypergeometric test) for ORF9B-binding partners.

(B) Molecular functions enriched in ORF9B interactors. Bar plot illustrates significantly enriched molecular functions (FDR<0.05, hypergeometric test) among ORF9B-associated proteins identified by APPLE-MS.

In (A) and (B), the length of each bar corresponds to the -$\log_{10}$(FDR) value, representing the statistical significance of enrichment.
